# Identification of a Novel Gene *argJ* involved in Arginine Biosynthesis Critical for Persister Formation in *Staphylococcus aureus*

**DOI:** 10.1101/114827

**Authors:** Rebecca Yee, Peng Cui, Tao Xu, Wanliang Shi, Jie Feng, Wenhong Zhang, Ying Zhang

## Abstract

*Staphylococcus aureus* can cause both acute and recurrent persistent infections such as peritonitis, endocarditis, abscess, osteomyelitis, and chronic wound infections. An effective treatment to eradicate the persistent disease is still lacking as the mechanisms of *S. aureus* persistence are poorly understood. In this study, we performed a comprehensive and unbiased high-throughput mutant screen using *S. aureus* USA300 and identified *argJ,* encoding an acetyltransferase in the arginine biosynthesis pathway, whose mutation produced a significant defect in persister formation in multiple drugs and stresses. Genetic complementation and arginine supplementation restored persistence in the ArgJ mutant. Quantitative real-time PCR analysis showed that the *arg* genes were over-expressed under drug stressed conditions and in stationary phase cultures. In addition, the ArgJ mutant had attenuated virulence in both *C. elegans* and mouse models of infection. Our studies identify a novel mechanism of persistence mediated by arginine metabolism in *S. aureus.* These findings will not only provide new insights about the mechanisms of *S. aureus* persistence but also offer novel therapeutic targets that may help to develop more effective treatment of persistent *S. aureus* infections.

## Introduction

Persisters are metabolically quiescent cells that are tolerant to antibiotics or stresses but can revert back to a growing state upon antibiotic stress removal and remain susceptible to the same antibiotic^1^. Persister cells are implicated in persistent infections^2^ and can cause relapse in various bacterial infections. While *S. aureus* often causes acute infections, it can also cause chronic recurrent infections such as peritonitis, endocarditis, osteomyelitis, wound and soft tissue infections, and infections from indwelling medical devices^3^.

Most studies on persister cell mechanisms have been conducted using *Escherichia coli* (*E. coli*) as a model organism. In *E.coli,* the toxin-antitoxin (TA) systems such as HipBA and MazF cause persister formation by inhibiting protein synthesis through phosphorylation of Glu-tRNA synthase and cleaving of mRNA, respectively ^4^. Unlike *E. coli,* it has been shown that *S. aureus* persister formation does not involve TA systems but is dependent on ATP production ^5^. Additionally, we have also previously shown that pathways involved in protein synthesis, efflux/transporter and metabolism and energy production are heavily involved in persister cell formation in *S. aureus* ^6^. We have shown that upon prolonged exposure to rifampicin, over one-hundred genes that play a role in rifampicin persistence were identified, where approximately one-third of the genes play a role in the metabolism of amino acids, lipids, vitamins, carbohydrates and purine biosynthesis ^6^.

To find core genes and pathways that play a role in persister cell formation with different drugs, we performed an unbiased high-throughput screen using gentamicin against a mutant library of *S. aureus* clinical isolate USA 300^7^. We identified the *argJ* gene as a core gene that plays an important role in persister formation under various antibiotics and stresses. We report for the first time the importance of ArgJ for *S. aureus* persister cell formation and also virulence and survival in *C. elegans* and mice.

## Results

### argJ is a novel persistence gene

To identify genes and pathways involved in persister formation, we performed a genome-wide screen using the saturated Nebraska transposon mutant library (NTML)^7^ to isolate mutants with defective persistence which is defined as a decrease in the number of persister cells relative to the parental strain USA300. We exposed stationary phase cultures of the mutant library to gentamicin (60 μg/ml, 10X MIC) (Fig. 1A). Over the course of six days, the screen identified the ArgJ mutant that failed to grow on tryptic soy agar (TSA) plates after gentamicin exposure. While it is known that the *argJ* gene is required for the arginine biosynthesis cycle^8 9^ and encodes an acetyltransferase that synthesizes N-acetylglutamate and ornithine (Fig. 1B), the importance of ArgJ in antibiotic and stress tolerance has not been explored.

**Figure 1.**
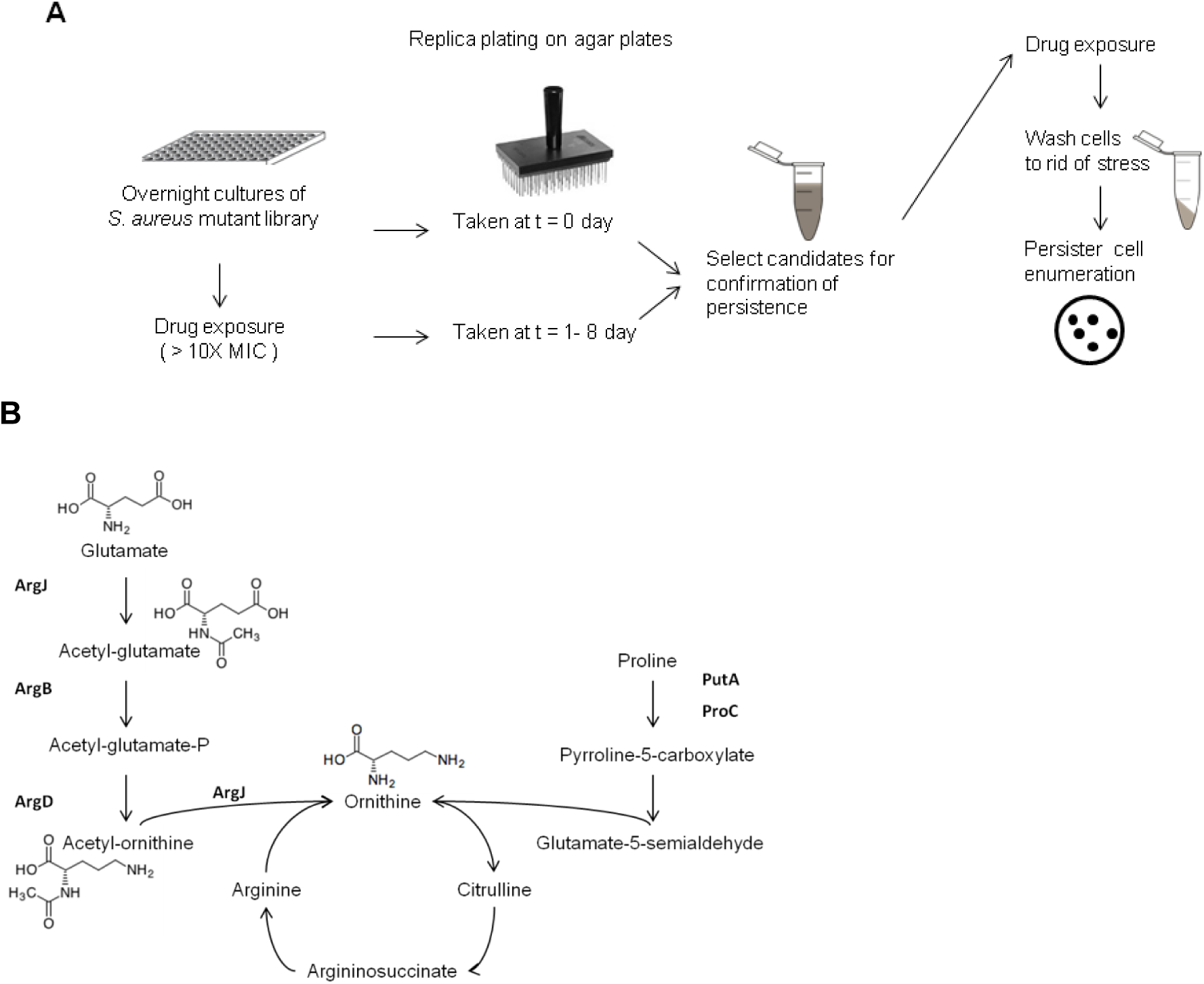
Summary of the genetic screen. (A) Work flow of genetic screen. (B) The arginine biosynthesis pathway in *S. aureus.*

To establish the importance of ArgJ in stress tolerance, we first excluded the possibility that altered growth dynamics is a confounding factor. Our growth curve study suggested that the ArgJ mutant and USA300 had similar growth patterns (Fig 2A) and similar colony forming unit per milliliter (CFU/ml) in normal growth medium even up to 8 days (Fig 2B). Next, to confirm that a mutation in ArgJ causes a defect in persister formation in stressed conditions, we performed a persister assay by exposing stationary phase cultures of the ArgJ mutant and USA300 control strain to different antibiotics and stresses. At different time points, the cells were washed and then enumerated for CFU. As early as day 2 post-gentamicin exposure, the ArgJ mutant harbored 1× 10^7^ CFU/ml as opposed to USA300 with 1×10^9^ CFU/ml. By day 6, the ArgJ mutant had 1 ×10^6^ CFU/ml compared to USA300 with 1 × 10^9^ CFU/ml, a significant difference of about 3-logs (Fig. 2C). Similarly, by day 6 of rifampicin exposure, there was also a significant three-log difference in CFU between the ArgJ mutant and USA300 (Fig. 2D).

**Figure 2.**
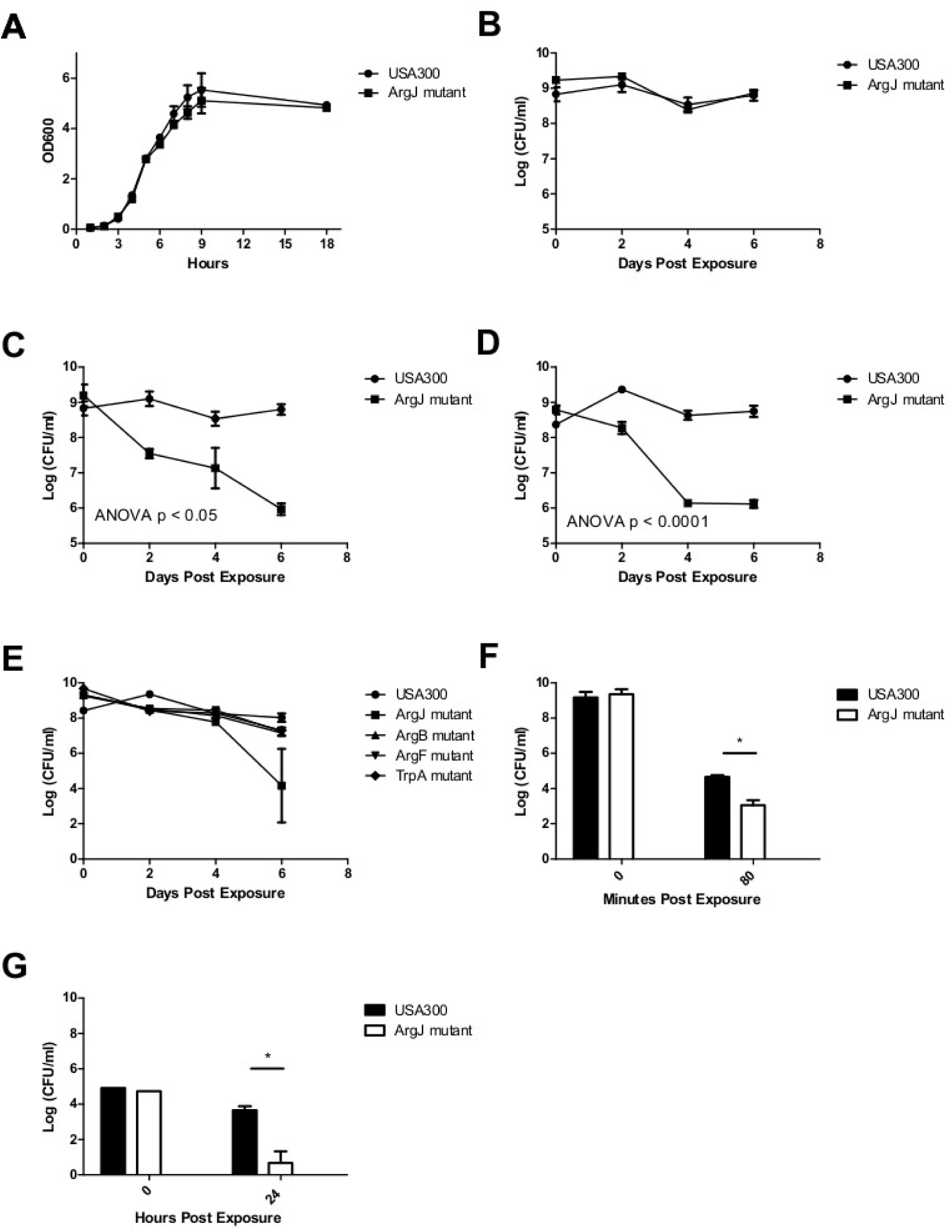
Phenotypes of parental strain USA300 and the ArgJ mutant. (A & B) Parental strain USA300 and the ArgJ mutant had no difference in growth dynamics. The ArgJ mutant showed defective persistence in both drugs (C) gentamicin (60 μg/ml, >10X MIC) and (D) rifampicin (2 μg/ml, > 10XMIC). (E) Strain with mutations in ArgB, ArgF, and TrpA did not show a defect in persistence. More killing was seen in the ArgJ mutant by (F) heat stress of 58°C and (G) low pH of 4.0. Data are representative of three independent experiments. * = p< 0.05 by two-way ANOVA or Student's t-test.

To confirm the specific role of ArgJ in persistence, we also measured the persister levels in mutants with mutations in other proteins of the Arg pathways (e.g. ArgB and ArgF) and TrpA, protein involved in tryptophan biosynthesis, as a control. Our results indicated that after gentamicin exposure of 6 days, there were no significant differences in CFU/ml among the ArgB, ArgF, TrpA mutants and the control strain USA300 (Fig. 2E). We, therefore, concluded that a mutation in ArgJ is specific in causing a defect in persister formation in *S. aureus.*

In our separate recent study, we showed that mutations in metabolic pathways in *S. aureus* have defective persistence to different antibiotics, low pH, and heat stress ^6^. To explore if ArgJ mediates persister cell formation in other stresses besides antibiotics, we subjected the ArgJ mutant to heat stress at 58°C. The difference in heat tolerance between the ArgJ mutant and USA300 was statistically significant. After 80 minutes, the ArgJ mutant had a mean of 1.1 × 10^3^ CFU/ml while USA300 had 4.6 × 10^4^ CFU/ml (Fig. 2F). Additionally, the ArgJ mutant was also significantly less tolerant to low pH (pH = 4) compared to USA300. Unlike the other stress exposures, the starting bacterial inoculum concentration for acid pH exposure was standardized to 1 × 10^5^ CFU/ml in order to prevent neutralization of the acid pH due to neutralization of acid pH by a high bacterial inoculum ^10^. Nonetheless, after 24 hours of exposure in a low pH environment, the ArgJ mutant had only about 4.6 CFU/ml left while the USA300 had about 4.4 × 10^3^ CFU/ml, indicating that the ArgJ mutant is more susceptible to low pH (Fig. 2G).

### Complementation of ArgJ mutant partially restored persistence phenotype

To confirm that the mutation in *argJ* is responsible for the defective persistence, we complemented the ArgJ mutant with the wildtype *argJ* gene from the USA300. We used an *S. aureus-E. coli* shuttle vector pRAB11 to insert the wildtype *argJ* gene back into the mutant ^11^. After 6 days of gentamicin exposure, the ArgJ mutant with an empty vector had 6.4 × 10^3^ CFU/ml whereas the ArgJ complemented strain and USA300 had 7.1 × 10^4^ CFU/ml and 2.0 × 10^7^ CFU/ml, respectively (Fig. 4A). Similarly, after 6 days of rifampicin exposure, the ArgJ mutant with an empty vector had 3.1 × 10^3^ CFU/ml whereas the ArgJ complemented strain had 5.0 × 10^6^ CFU/ml, only one-log fold less than USA300 with 1.7 × 10^7^ CFU/ml. These findings indicate that a genetic mutation in ArgJ confers a defect in persister formation.

### Arginine biosynthesis via the Arg pathway is important for persistence

To determine if the arginine pathway is important for persistence, we supplemented L-arginine into TSB growth medium. The ArgJ mutant grown without any L-arginine supplementation had 1.2 × 10^4^ CFU/ml under gentamicin stress. However, when L-arginine (30 mM) was supplemented into the growth medium of the ArgJ mutant, the amount of cells on day 6 post-gentamicin exposure was 6.0 × 10^6^ CFU/ml, which is similar to the parent strain USA300 with 2.8 × 10^6^ CFU/ml (Fig. 3C), indicating that L-arginine supplementation complemented the defect in persistence in the ArgJ mutant. Because arginine is a positively-charged amino acid, we wanted to confirm that persistence restoration is specific to arginine and not achieved by other positively-charged amino acids. Thus, we supplemented the growth medium with two positively-charged amino acids L-histidine and L-lysine. Our results suggest that histidine and lysine did not restore the persistence of the ArgJ mutant indicating that L-arginine is specifically important for persistence (Fig. 3C).

**Figure 3.**
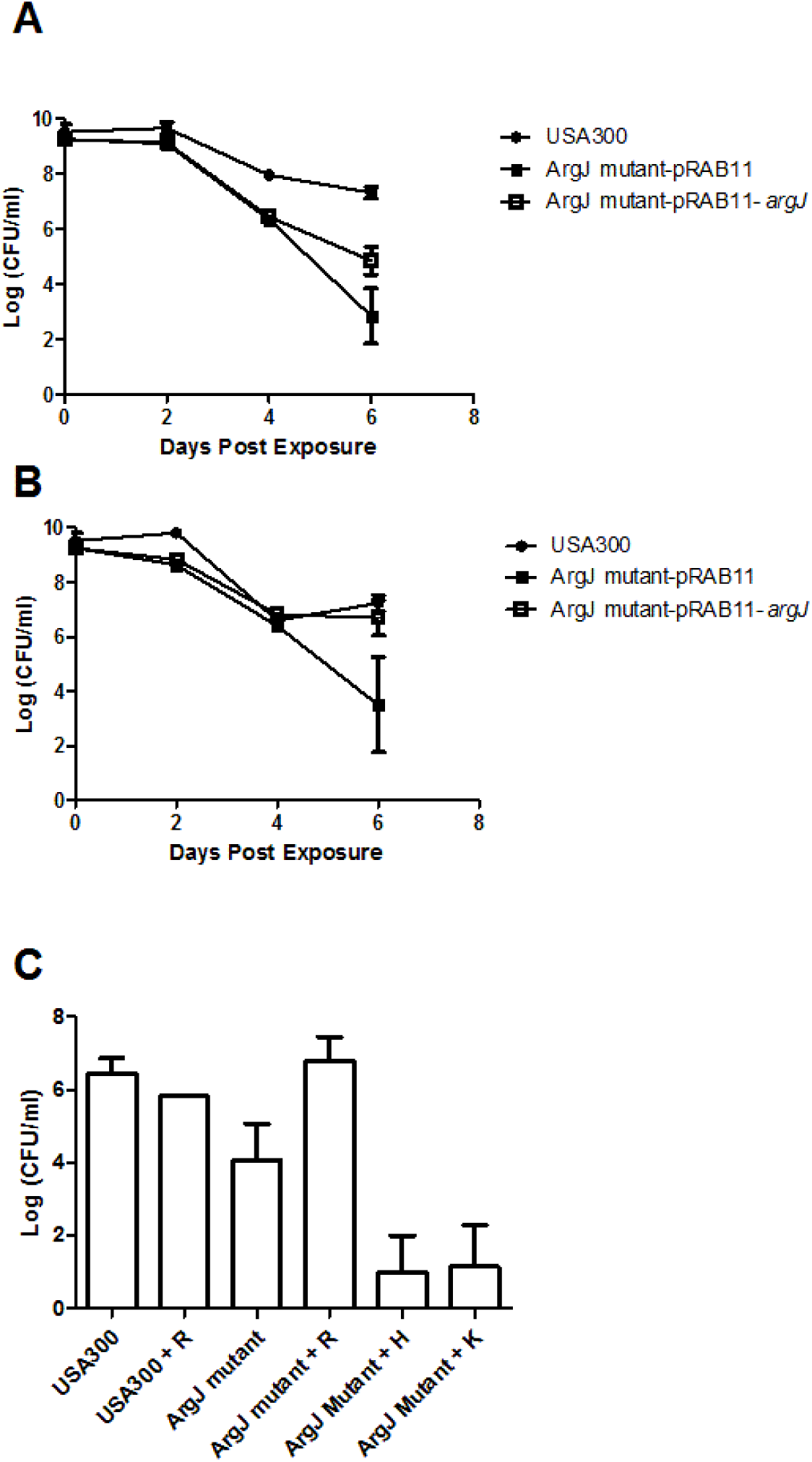
Persistence in antibiotics can be restored in the ArgJ mutant. (A) The ArgJ mutant was transformed with the wildtype *argJ* gene using shuttle vector pRAB11. Upon 6 day exposure of (A) gentamicin and (B) rifampicin, the ArgJ mutant complemented with *argJ* gene showed partial restoration of persistence compared to the empty vector. (C) The supplementation of L-arginine (30 mM) can restore gentamicin persistence. The effect of amino acid supplementation to rescue ArgJ's persistence phenotype is specific to the amino acid arginine. Amino acids with similar chemical properties to arginine (eg. histidine and lysine) did not restore persistence. Data are representative of three independent experiments.

### Activity of arginine pathway genes in relation to persistence

*S. aureus* has the ability to synthesize arginine using secondary carbon sources such as glutamate (via the Arg pathway) or proline (via PutA and ProC) (Fig. 1B)^12^. It has been suggested that arginine production under normal growth conditions is mainly due to the proline precursor pathway^12^. However, the activity of the Arg pathway under stress conditions such as stationary phase and antibiotic exposure is unknown. To evaluate if the Arg pathway is induced under stress conditions, we performed qRT-PCR to compare the levels of gene expression of genes from the Arg pathway (*argCG*) versus genes involved in arginine synthesis from a proline precursor (*proC* and *putA*). Since the ArgJ mutant showed defective persistence (Fig. 2) compared to USA300, we compared the gene expression fold-change of USA300 and the ArgJ mutant. Genes *argC* and *argG* were at least 2-fold more over-expressed in USA300 than the ArgJ mutant (Fig. 4A) in stationary phase (when persister cells enrich due to limiting nutrients) compared to log phase (when growing cells are heavily populated). Under gentamicin treatment, we also observed the same genes, *argC* and a*rgG*, having at least 2-fold higher expression in USA300 than in the ArgJ mutant (Fig. 4B). Collectively, our data suggest that arginine production through the ArgJ pathway is more expressed in stationary and drug-treated cells than log phase cells and untreated cells, respectively.

**Figure 4.**
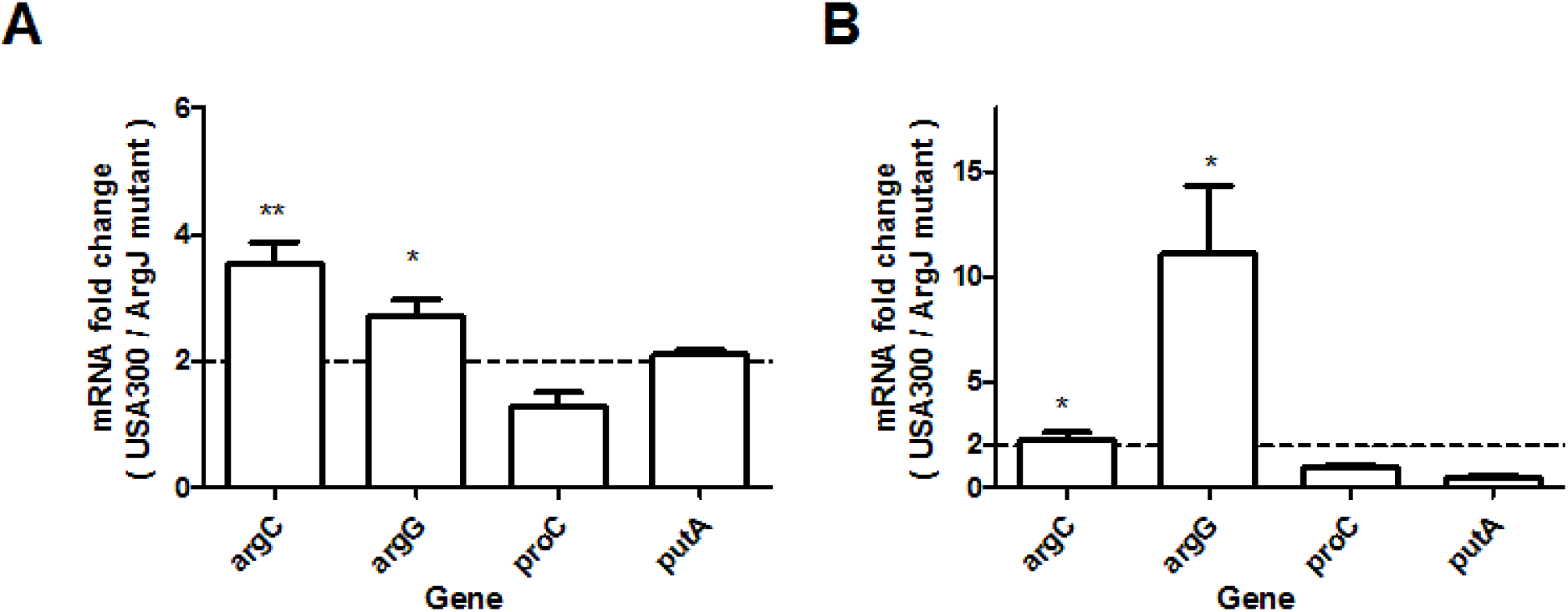
Arginine biosynthesis through the Arg pathway plays a role in persistence. In stationary phase (B) or gentamicin-treated (C) cultures, there is at least 2-fold more expression of Arg pathway genes *argC* and *argG* in USA300 compared to the ArgJ mutant. Differences among the expression of the *arg* genes and both the expression of *proC* and *putA* are also statistically significant. Data are representative of three independent experiments. ** = p < 0.005, * = p < 0.05 by Student's t-test.

### ArgJ plays a role in virulence in vivo

There is a general lack of research establishing the relationship between the mechanisms of persister cell formation and virulence in different bacterial pathogens including *S. aureus*^2^. Given that our results suggest the importance of ArgJ in persistence, we decided to explore the role of ArgJ in virulence. In order to test for virulence, we used a nematode *C. elegans* model, an accepted model for bacterial pathogenesis research^13^. To test the hypothesis that the ArgJ mutant has attenuated virulence, we examined the survival of *C. elegans* after *S. aureus* infection. Our results showed that *C. elegans* killing was highly attenuated after infection with the ArgJ mutant as opposed to USA300. The first death caused by USA300 was observed at day 4 as opposed to day 6 for the ArgJ mutant. By day 8, there was a 76% survival in *C. elegans* exposed to the ArgJ mutant as opposed to the 56% survival in worms exposed to USA300 (Fig. 5A). No change in survival was seen in worms exposed to nonpathogenic *E. coli* strain.

To further confirm the role of ArgJ in virulence, we then utilized an *S. aureus* peritonitis mouse model ^14^ Briefly, Swiss-Webster mice were infected by intraperitoneal injection with 7 × 10^7^ CFU of the ArgJ mutant, the ArgJ complemented strain and USA300. After 3 day post-infection, the CFU counts in the spleens and kidneys were enumerated. Our results indicate that mice infected with the ArgJ mutant harbored an average of 5 CFU/g of spleen while mice infected with USA300 and the ArgJ complemented strain had 3.5 × 10^4^ CFU/g of spleen and 1.6 × 10^3^ CFU/g of the spleen, respectively (Fig 5B). Similarly, all mice infected with the ArgJ mutant resulted in no bacteria in the kidney while mice infected with USA300 and the ArgJ complemented strain had 1.0 × 10^4^ CFU/g and 3.6 × 10^3^ CFU/g of the kidney, respectively (Fig 5C), indicating that an ArgJ mutation caused statistically significant attenuation of virulence in mice.

**Figure 5.**
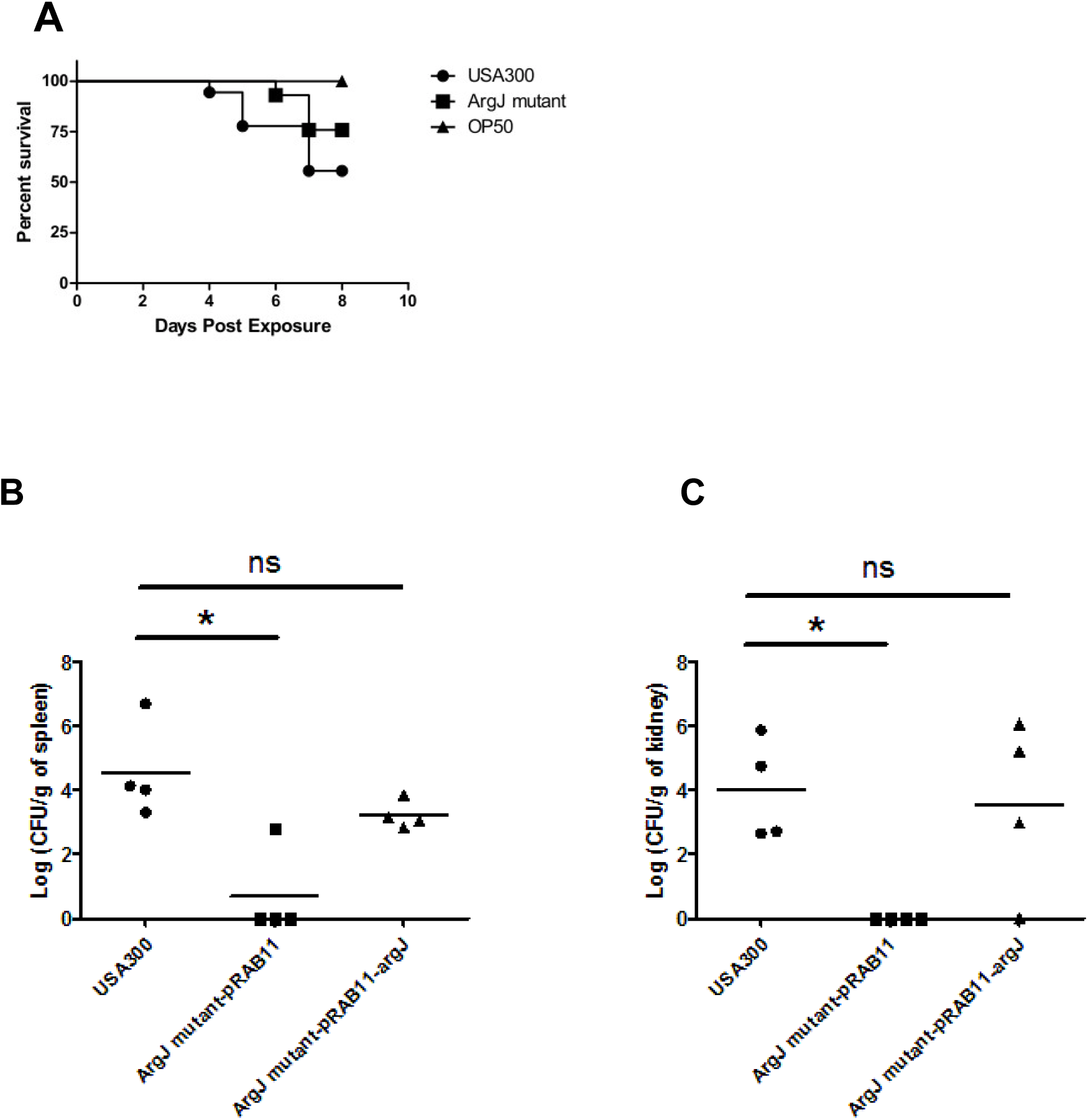
The ArgJ mutant has attenuated virulence in-vivo. (A) The ArgJ mutant showed attenuated killing of *C. elegans* compared to parent strain USA300. Three days post-infection, mice (n=4) infected with the ArgJ mutant had a lower bacterial load in both the (B) spleens and (C) kidneys. Genetic complementation with the wildtype *argJ* gene restores virulence in the ArgJ mutant. (* = p < 0.05 by Student's t-test).

## Discussion

While there is renewed interest in persister biology recently, the molecular mechanisms of persistence are largely derived from the model organism *E. coli* and the mechanisms crucial to *S. aureus* persister formation remain poorly understood. After we performed a comprehensive genetic screen to identify persister genes and pathways in the clinically relevant strain USA300, we identified ArgJ as important for persistence to multiple drugs and also stresses. To our knowledge, this is the first study that provided insights into the physiological impact of ArgJ and its related Arg pathway in stress tolerance and persistence in bacterial systems.

Supplementation of arginine but not other amino acids into the growth media conferred increased persistence of the ArgJ mutant to gentamicin. Our data suggest that arginine increases tolerance and formation of persister cells and that ArgJ regulates persistence in *S. aureus* is further supported by our qRT-PCR results. It is important to note that under normal growth conditions, our results are consistent with previous findings that suggest arginine synthesis from the proline precursor pathway (using PutA and ProC) is the canonical pathway (Fig. 2B)^12^. However, the ArgJ-mediated Arg pathway appears to play a role in stationary phase and during stress conditions (Fig. 5).

While the molecular mechanisms by which the arginine biosynthesis pathway mediates persistence remains to be determined, there are several possibilities. One proposed mechanism of ArgJ-mediated persistence is through the direct generation of arginine. The bacterial cell can then catabolize the arginine through the arginine deiminase pathway to synthesize ammonia which mitigates against hydroxyl radicals produced during antibiotic action that promote cell death ^15^. Persister cells are known to have increased capacity to deal with reactive oxygen species^2^. Additionally, the downstream products of arginine production such as ornithine and polyamines are shown to increase the cell's fitness and survival ^16^. Polyamines also modulate the translation and expression of key proteins in biofilms, an exopolysaccharide structure that contains persister cells^17^. Because persisters are non-growing cells with decreased energy state^2^, we speculate that an ArgJ mutation causes defective persistence because the mutant bacteria need to resort to a more energy unfavorable pathway to produce arginine. In *S. aureus,* ArgJ is preferred over ArgE, which produces the same end products of ornithine and acetate, due to favorable energy kinetics ^9^. Thus, under growth-limiting conditions where the cells are more energetically inactive, ArgJ may be preferentially expressed to facilitate persister survival. Hence, the altered cellular energetic state could impede the cells to reach dormancy and allow the ArgJ mutant to be killed more easily by antibiotics and stresses.

ArgJ is a bifunctional enzyme involved in de novo as well as recycling pathway for arginine biosynthesis (Fig 1B). The role of ArgJ in the de novo pathway of arginine synthesis is to catalyze the first step of the linear arginine production, synthesizing N-acetylglutamate from glutamate and acetyl-CoA as the acetyl donor. In the recycling pathway, ArgJ helps generate ornithine by trans-acetylation of the acetyl group from N(2)-acetylornithine to glutamate. Our finding that mutations in de novo pathway genes (*argB, argF*) did not cause a defect in persistence (Fig. 3) indicates that the de novo arginine biosynthesis pathway is not important for persistence in *S. aureus* but rather the recycling function of ArgJ may be important for persistence. Further biochemical and genetic studies such as site-directed mutagenesis on the binding and active sites of the ArgJ protein to separate the bifunctional activity of the protein are needed to confirm the importance of the recycling pathway in persistence.

In addition to ArgJ, 6 of the genes that we have identified playing a role in gentamicin persistence were also transferases (*miaA, trmB, SAUSA300_ 0689, 1111, 1669, 2232*) (data unpublished). Acetyl-transferases can help drive bistable gene expression and changes in DNA and protein modifications in tolerating stress and adapting to environmental changes^2,18^. Studies have shown that acetylated proteins in prokaryotes play a role in stress response through chemotaxis and cell cycle control ^19^. Protein homology analyses suggest that the binding site of acetyltransferase SAUSA300_2232, also identified as a gene important for persister formation, has a similar amino acid sequence to ArgJ. More studies will be needed to explore the role of these transferases in epigenetic control of persistence in *S. aureus.*

Our complementation studies showed that the *argJ* gene is important for maintaining persistence in *S. aureus.* However, the complemented ArgJ mutant achieved partial restoration of persistence. In fact, in virtually all *S. aureus* complementation studies, none could achieve full complementation. *S. aureus* possesses different restriction modification systems that may destroy exogenous plasmids,^20^ has endonucleases targeting specific sequences^21^ and can methylate exogenous DNA ^22^ causing inactivation of exogenous DNA. These could serve as possible explanations for the partial complementation of ArgJ mutant in this study.

While this study revealed novel insights into the mechanisms of *S. aureus* persister formation, the results from the screen is dependent on the conditions of our assays^2^. Persisters can be affected by variables pertaining to the drugs administered such as the drug concentrations, drug exposure time, inoculum size and the age of the culture ^10^. However, the reported persister genes in this study (Tqable S1) have been identified twice from two independent screens with both rifampicin and gentamicin and can thus be considered reproducible core genes involved in persister formation in *S. arueus.*

In conclusion, we identified a comprehensive list of genes and pathways that play a role in establishing persistence in *S. aureus.* For the first time, we identified a novel mechanism of persistence in *S.* aureusmediated by ArgJ in maintaining persistence to different antibiotics and stresses and also virulence in-vivo. For *S. aureus,* tackling persistence may be a solution to reducing the rate of drug resistance. Our findings not only improve our understanding of mechanisms of persistence but also provide insights into novel therapeutic targets for developing new and more effective drugs that eradicate persistent *S. aureus* infections.

## Experimental Procedures

### Culture media, chemicals, and antibiotics

*S. aureus* strains were cultivated in tryptic soy broth (TSB) and tryptic soy agar (TSA) and *E. coli* strains were cultivated in Luria-Bertani (LB) broth or agar at 37°C with the appropriate antibiotics. Citric acid monohydrate, the antibiotics ampicillin, chloramphenicol, rifampicin, gentamicin, erythromycin and amino acids L-Arginine, L-Histidine, L-Lysine were obtained from Sigma-Aldrich Co. Stock solutions were prepared and sterilized through filtration or autoclaved, if necessary, and used at indicated concentrations.

### Library screen to identify mutants with defective persistence

The Nebraska Transposon mutant library (NTML) that consists of 1,920 mutants of *S. aureus* USA300 was kindly provided by BEI Resources^7^. The mutants of the library were grown in TSB containing erythromycin (50 μg/ml), the antibiotic selective marker of the mutants, at 37°C in 384-well plates. Gentamicin (60 μg/ml) was added to overnight stationary phase cultures in the wells. The plates were incubated at 37°C and the library was replica transferred to TSA plates to score for mutants that failed to grow over the course of at least 6 days.

### Persister assays to measure susceptibility to various antibiotics and stresses

Overnight stationary phase cultures were exposed to selected drugs or stresses and colony forming units per milliliter (CFU/ml) were measured through serial dilutions and plating onto TSA plates. The antibiotic exposure was carried out over the course of 6 days at 37°C as previously described ^10^. To measure the susceptibilities in low pH stress, the overnight culture was diluted 1:100 and incubated in buffered acid solution with pH = 4 at 37°C. To measure the susceptibility to heat, undiluted overnight cultures were placed in a 58°C water bath. At different time points, 100 μl of bacterial suspension was removed and washed in 1X PBS and enumerated for CFU/ml. For amino acid supplementation, overnight bacterial cultures were refreshed 1:100 into TSB with a supplementation of the indicated amino acids to the growth media and grown for 16 hours at 37°C before use. This step was repeated again before persister assays were performed.

### Complementation of ArgJ mutant

The wildtype *argJ* gene from *S. aureus* USA300 was amplified by PCR. The primers contained restriction sites KpnI and EcoRI. The forward primer used had the sequence 5' GCAGGTACCATGAAACATCAAGAAACGAC 3' and the reverse primer used had the sequence 5' GCCGAATTCTTATGTTCGATATGATGCGTT 3'. The PCR parameters used were as follows: 94°C for 15 min, followed by 35 cycles of 94°C for 30 s, 55°C for 30 s, and 72°C for 2 min, and a final extension at 72°C for 10 min. The PCR products were digested with KpnI and EcoRI and cloned into *S. aureus-E. coli* shuttle vector pRAB11 ^11^. Ligation mixtures were transformed into chemically competent *E. coli* DH5α cells (Invitrogen) and spread onto LB agar plates containing ampicillin (100 μg/ml) and grown overnight at 37°C. Upon confirmation of the transformants by DNA sequencing, plasmid DNA was isolated using Lysostaphin (Sigma-Aldrich) lysis (2mg/ml) followed by purification with QIAprep Spin Miniprep Kit. The plasmid was introduced into *S. aureus* RN4220^23^ by electroporation (voltage = 2.5 kV, resistance = 100 Ω, capacity = 25 μF) using MicroPulser Electroporation Apparatus (Bio-Rad) followed by plating onto TSA plates containing chloramphenicol (10 μg/ml) and incubation overnight at 37°C. To induce for ArgJ expression, bacterial cultures were grown in TSB containing chloramphenicol overnight then refreshed 1:100 into TSB only. When the cells reached OD600 of 0.5, the cells were induced with anhydrotetracycline (25 ng/ml) overnight with shaking at 220 rpm in 37°C. Overnight cells were washed twice with 1X PBS and resuspended in MOPS buffer to perform persister assays as described above.

### RNA preparation and real-time PCR (qRT-PCR)

Samples were prepared for RNA extraction based on the instructions described in the "Enzymatic Lysis and Proteinase K Digestion of Bacteria" protocol of the RNAprotect Bacteria Reagent Handbook with the addition of incubation with lysostaphin before purification using the RNeasy mini kit (Qiagen). cDNA was synthesized from 1 μg of RNA with random primers using QuantiTech Reverse Transcription Kit (Qiagen). Quantitative RT-PCR (qRT-PCR) was performed in a 20 μl reaction mixture using SYBR Green PCR Master Mix (Life Technologies) and 0.2 μM (each) of gene-specific primers (Table S2). Amplification and detection of specific products were performed using StepOnePlus Real-Time PCR Systems (Applied Biosystems). The PCR parameters used were as follows: 95°C for 10 min, followed by 40 cycles of 95°C for 15 s and 60°C for 1 min. Relative gene expression levels were calculated using the comparative threshold cycle (C_T_) method (2^−∆∆CT^ method) with 16s rRNA as the internal control gene for normalization of gene expression to basal levels.

### Nematode-killing assay

*S. aureus* nematode-killing assay was performed as described^13^. Briefly, *S. aureus* strains were grown overnight at 30°C in TSB containing the appropriate antibiotics as needed. One spot of overnight *S. aureus* culture (70 μl) was dropped onto nematode growth agar containing 5- Fluoro-2'-deoxyuridine (100 μM). The prepared plates were incubated at 37°C overnight and then allowed to equilibrate to room temperature (20°C) for 60 minutes before being seeded with *C. elegans* N2 Bristol worms *(Caenorhabditis* Genetics Center). The worms were synchronized to the same growth stage by treatment with alkaline hypochlorite solution as described^24^. Worms of the adult stage were recovered in 15 ml tubes with M9 buffer. The worms were washed twice to remove the residual bacteria in their diet by centrifugation at 1500 rpm for 2 minutes at room temperature in a table top centrifuge. Bleaching solution with 5% hypochlorite was then added and incubated with the worms for 9 minutes to lyse the adult stages but keeping the eggs intact. The lysing reaction was stopped when M9 buffer was added. Bleach was removed by centrifugation at 1500 rpm for 1 minute followed by three more washes with M9 buffer. To induce hatching of eggs, M9 buffer was added to the pellet and incubated at 20°C with gentle agitation and proper aeration. After 24 hours, worms were pelleted with a 2-minute spin at 1500 rpm at room temperature and seeded onto OP50 seeded plates. L4 stage worms were obtained after 48 hours at 20°C. In each assay, 10-20 L4-stage nematodes were added to each plate and each assay was carried out at least twice. The plates were incubated at 20°C and scored for live and dead worms every 24 hours. A worm was considered dead when it failed to respond to touch.

### Mouse intraperitoneal challenge

Overnight cultures of *S. aureus* were subcultured into fresh TSB (1:100) and grown for 2.5 hours with shaking (220 rpm) at 37°C. The cells were washed with 1X PBS. Adult (7-8 weeks old) female Swiss-Webster mice (Charles River Laboratories) were infected via intraperitoneal injection with an inoculum size of 7 × 10^7^ CFU. Mice were housed in cages under standard BSL-2 housing conditions. Mice infected after 3 days were euthanized and spleens and kidneys were homogenized for CFU enumeration.

## Acknowledgements

We gratefully acknowledge BEI Resources for the provision of the NTML mutant library. We thank Jiou Wang and his laboratory for the provision of *C. elegans* and discussions regarding the design of nematode killing assays. RY was supported by NIH training grant T32 AI007417. YZ was supported by NIH grants AI099512 and AI108535.

## Author Contributions

R.Y., P.C., J.F., W.S., and W.Z. contributed to the design of the study. R.Y., P.C., J.F., and Y.Z. helped with the acquisition, analysis and interpretation of the data. R.Y. and Y.Z. wrote the paper.

---

**Table S1.**
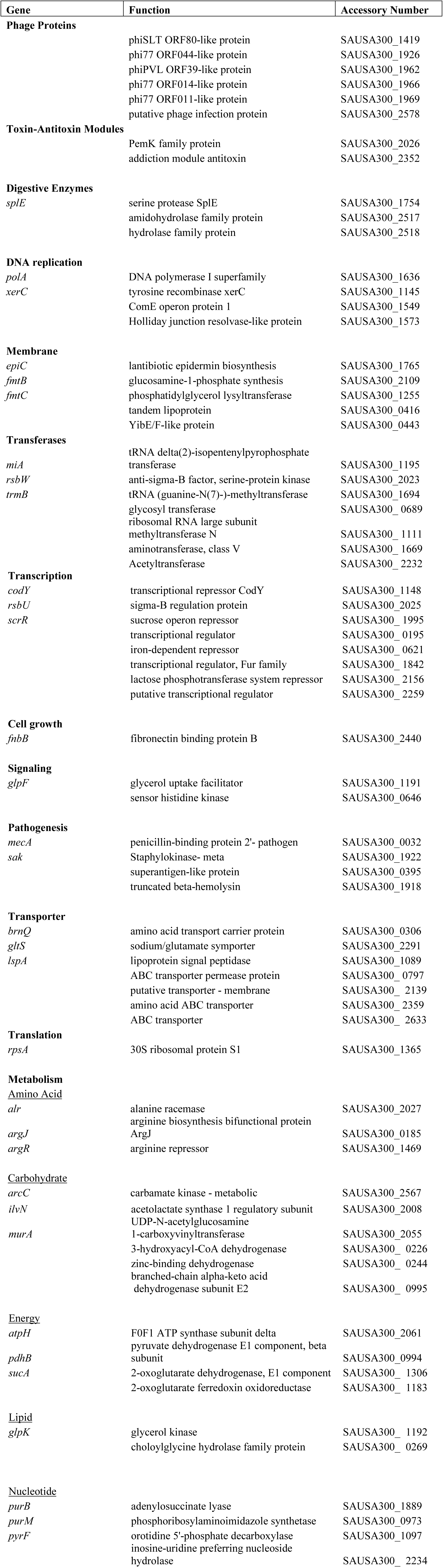
Genes that play a role in gentamicin persistence

**Table S2.**
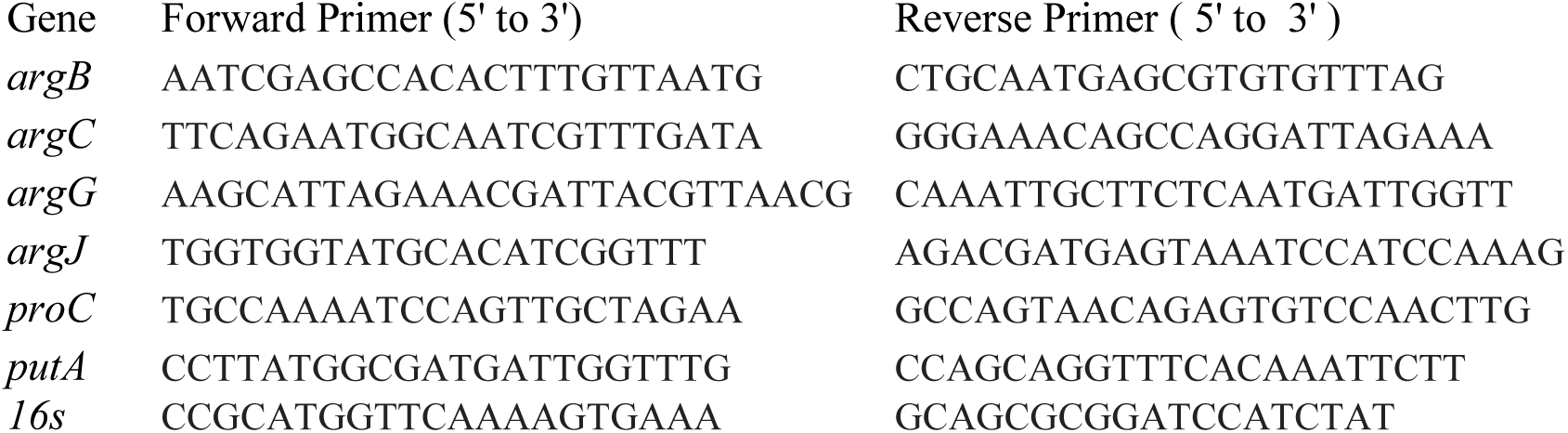
Oligonucleotide primers used for qRT-PCR

